# Global diversity and distribution of coral-associated protists

**DOI:** 10.64898/2026.01.22.700988

**Authors:** Javier del Campo, Anthony M. Bonacolta, Bradley A. Weiler, Ben Knowles, Amy Apprill, Michael D. Fox, Kevin C. Wakeman, Mark J. A. Vermeij, Forest Rohwer, Patrick J. Keeling

## Abstract

Coral reefs are critical ecosystems and biodiversity hotspots that provide ecological stability and essential services to coastal communities. The coral holobiont, a complex symbiotic system composed of the coral animal and a diverse array of associated microbes, plays a central role in coral health and resilience. While Symbiodiniaceae and bacterial symbionts have been extensively studied, much less is known about the diversity and function of microbial eukaryotes such as protists and fungi. These organisms are increasingly recognized as important, yet remain vastly underexplored. Here, we present the first global survey of the coral-associated eukaryome using an anti-metazoan 18S rRNA primer set to bypass host DNA amplification. Our dataset includes corals and related anthozoans from the Caribbean Sea, the Red Sea, and several locations in the Pacific Ocean, spanning a broad taxonomic and geographic range, and includes both healthy and diseased specimens. They reveal a eukaryome that is not only more diverse than that of global coastal waters but also surpasses the diversity of the well-studied coral bacterial microbiome. We recover diverse microbial eukaryotic communities, including Symbiodiniaceae, other known symbionts, potential pathogens, and previously uncharacterized lineages. These results reveal consistent patterns across coral groups and geographic regions. This study provides the most comprehensive taxonomic overview of coral-associated microbial eukaryotes to date, offering new insights into their roles within the holobiont. Our findings highlight the ecological significance of microbial eukaryotes and underscore the importance of incorporating them into broader coral reef research and conservation strategies.

## Background

Corals are among the most important architects of marine ecosystems, shaping the seafloor from shallow coastal environments to deeper ocean regions. Coral reefs serve as biodiversity hotspots, supporting many marine species that rely on these habitats as their ecological niches[1]. Beyond their environmental importance, coral reefs provide essential ecosystem services. They protect shorelines from storm surges, serve as vital resources for fisheries, and attract millions of tourists annually, supporting local economies [2, 3].

However, coral reefs face unprecedented threats from climate change, including marine heatwaves and emerging diseases[4, 5]. This underscores the urgent need to expand our understanding of corals and develop effective strategies to mitigate these impacts. Still, major gaps remain in our understanding of even some of the most fundamental aspects of this complex system, and one of these is the diversity of microbial eukaryotes associated with corals.

Corals are a multisymbiotic system, collectively known as the coral holobiont [6]. The holobiont comprises the coral animal host and a diverse community of microorganisms, including bacteria, archaea, viruses, fungi, algae, and other protists. The interactions among these partners are fundamental to the health and survival of the coral host. The best-known symbiosis is between the coral holobiont and dinoflagellate algae, the Symbiodiniaceae [7]. These phototrophic algae reside in coral gastrodermal cells, supplying the host with photosynthates that constitute a significant portion of its daily carbon requirements [8, 9].

Symbiodiniaceae are highly sensitive to temperature changes and, during marine heatwaves, are expelled from the coral by a phenomenon known as bleaching, leaving the coral white in appearance and vulnerable to starvation [10]. If temperatures normalize, the coral may recover the algae; if not, the prolonged absence of this critical symbiont often leads to coral mortality. Bleaching events are becoming increasingly frequent and are responsible for the decline of a significant proportion of global coral cover.

Other microbial associates also play vital roles in the coral holobiont. Bacteria contribute to nutrient cycling, defense against pathogens, and overall holobiont stability [11]. Viruses, though less understood, are believed to influence microbial dynamics within the holobiont [12]. Unicellular microbial eukaryotes, such as fungi and protists, are also known to be present in corals, but our knowledge about these microbes remains much more limited [13]. Certain fungi, green algae, ciliates, labyrinthulids, and apicomplexans have been relatively well studied in coral, either as photosynthetic symbionts or agents of disease. The fungus *Aspergillus sydowii*, for example, causes disease in Caribbean sea fan corals (*Gorgonia spp.*) [14, 15]. Other fungi, particularly boring endolithic species, are linked to coral skeleton degradation [16]. The endolithic alga *Ostreobium querkettii* lives within the coral skeleton, sometimes at concentrations so high that a green ring is visible just below the living cnidarian [17]. Whether *Ostreobium* is a beneficial symbiont or parasite of the coral has been debated [18]. Still, given that it absorbs light at a wavelength not used by Symbiodiniaceae and provides photosynthates to the coral, it seems likely to be beneficial in most conditions [19]. Among heterotrophic protists, two types of ciliates are often associated with corals: the folliculinids [20] and the scuticociliates [21]. *Halofolliculina corallasia* was the first protozoan pathogen identified in corals, responsible for skeletal eroding band syndrome [22]. Labyrinthulomycetes are heterotrophic stramenopiles found on substrates in diverse habitats, including algal surfaces, mangrove leaves, seagrass, mollusks, and also coral mucus [23]. Although up to five Labyrinthulomycetes species have been isolated from corals [24, 25], and diverse members have also been reported to be widely associated with the coral *Fungia sp*., the nature of their interactions has not been elucidated [25, 26]. Apicomplexans are a large group of parasitic protists that cause devastating diseases such as malaria. These parasites evolved from photosynthetic ancestors, and photosynthetic sisters of the apicomplexans, the chromerids *Chromera velia* and *Vitrella brassicaformis*, were isolated from coral reefs [27, 28]. Until recently, a single apicomplexan species, *Gemmocystis cylindrus* [29], was reported from coral. However, metabarcoding studies and follow-up identification revealed a diverse group of apicomplexans called corallicolids infecting corals and related anthozoans [30–33].

Prior studies of microbial eukaryotes in corals relied heavily on microscopy and culture-based methods. While valuable, these approaches are labor-intensive and low-throughput. Advances in molecular techniques, such as high-throughput metabarcoding, have revolutionized the characterization of microbial communities in corals, particularly bacteria [34]. However, the application of these methods to microbial eukaryotes has been hindered by the overlap of genetic markers, such as the 18S rRNA gene, between the coral host and its symbionts. Recently, targeted primer sets have been developed to bypass host amplification, enabling more comprehensive studies of the coral-associated eukaryome [35]. These tools have already revealed the widespread presence of apicomplexans in Caribbean corals [31] or the abundant presence of Ichthyodinida (previously MALV – I or Dinoflagellates group I [36]) in hard and soft corals [37, 38] and hold promise for expanding our understanding of other eukaryotic symbionts.

Parasite diversity is often assumed to decline with ecosystem degradation, because many parasitic taxa rely on complex trophic webs and multiple hosts to complete their life cycles. Accordingly, we would expect pristine reefs to support a richer and more stable community of parasitic protists, while degraded reefs, with disrupted food webs, should host fewer lineages and a higher prevalence of opportunists. Our global survey of the coral eukaryome provides a unique opportunity to test this expectation by comparing parasite-associated groups such as apicomplexans, ciliates, and Ichthyodinida across reefs spanning gradients of environmental conditions.

Because corals are among the oldest extant animal lineages, with origins in the Precambrian, it is reasonable to expect that they have hosted microbial eukaryotes throughout their evolutionary history. We would therefore anticipate finding evidence of both deeply conserved protist associations dating back hundreds of millions of years, as well as more dynamic, recurrent colonizations by protist lineages that adapt to coral habitats over time. In this study, we conducted a global survey of the coral eukaryome using this anti-metazoan 18S rRNA approach, sampling corals and other anthozoans from diverse regions, including the Caribbean, the Red Sea, and the East, Central, and North Pacific Ocean. Our dataset encompasses many coral groups, including robust and complex scleractinians, black corals, octocorals, other anthozoans like sea anemones, zoanthids, corallimorphs, and related cnidarians such as hydrozoans and jellyfish, as well as both healthy and diseased specimens. This work provides the most comprehensive view of the coral eukaryome to date, filling critical knowledge gaps and offering insights into coral biology. Our findings deepen our understanding of coral-microbe interactions, providing fundamental knowledge needed to develop approaches to protect corals from decline.

## Methods

### Sampling

Corals were collected from several locations in Curaçao between April 19 and 21, 2015, and in Okinawa (Japan) between April 24 and 26, 2015. Whole samples, including skeleton and tissue, were rinsed using distilled water and homogenized using a mortar and pestle. DNA was extracted using the RNA Power Soil Total RNA Isolation Kit, coupled with the DNA Elution Accessory Kit (MoBio), for the Curaçao samples, and the DNeasy Blood &

Tissue Kit (Qiagen) for the Japan samples. DNA concentration was quantified on a Qubit 2.0 Fluorometer (Thermo Fisher Scientific Inc.). Samples from the Southern [39, 40] and Northern Line Islands [41], as well as the Red Sea [42] have been described previously in the cited manuscripts. A complete list of samples is available in Table 1.

### 18S rRNA amplification and Illumina sequencing

The V4 region of the eukaryotic 18S rRNA gene was amplified using a nested PCR approach validated in del Campo et al. 2019 [35]. Briefly, we first amplified the samples with the UNonMet primers to reduce the metazoan signal present [35], and then re-amplified these products using universal primers described by Comeau et al. [43]. Eukaryotic microbiome amplicons were prepared using PCR with high-fidelity Phusion polymerase (Thermo Fisher Scientific Inc.), using primers that target the V4 region of the 18S rRNA gene, but which exclude metazoan sequences (UNonMetF 5’-GTGCCAGCAGCCGCG-3’, UNonMetR 5’-TTTAAGTTTCAGCCTTGCG-3’) [44]. PCR was performed using the following protocol: 30s at 98°C, followed by 35 cycles each consisting of 10 s at 98 °C, 30 s at 51.1°C, and 1 min at 72°C, ending with 5 min at 72°C. PCR products were visually inspected for successful amplification using gel electrophoresis with 1% agarose gels. PCR products were then cleaned using the QIAquick PCR Purification Kit (Qiagen) and quantified on a Qubit 2.0 Fluorometer. Amplicon sequencing was performed by the Integrated Microbiome Resource facility at the Centre for Comparative Genomics and Evolutionary Bioinformatics at Dalhousie University. PCR amplification from template DNA was performed in duplicate using high-fidelity Phusion polymerase. A single round of PCR was done using “fusion primers” (Illumina adaptors + indices + specific regions) targeting the V4 region of the eukaryotic 18S rRNA gene (primer set E572F + E1009R [43]); (∼440 bp fragment) with multiplexing. PCR products were verified visually by running a high-throughput Invitrogen 96-well E-gel. The duplicate amplicons from the same samples were pooled in one plate, then cleaned-up and normalized using the high-throughput Invitrogen SequalPrep 96-well Plate Kit. The samples were then pooled to create a single library, which was quantified fluorometrically before sequencing on an Illumina MiSeq using a 300-bp paired-end read design. The 18S rRNA amplicon reads are deposited in the NCBI Sequence Read Archive (PRJNA482746 and PRJNA1373245).

### Bioinformatics analysis

Primers were removed from reads using Cutadapt v3.1 [45]. The trimmed reads were then processed in R using DADA2 [46]. Forward reads were truncated at 250 bp, and reverse reads were truncated at 190 bp. Reads were then denoised and joined into amplicon sequence variants (ASVs). Chimeras were removed using the ‘removeBimeraDenovo’ command with the “consensus” option. Taxonomy was assigned to ASVs using PR2 v4.13.0 [47]. Sequence tables, taxonomy tables, the phylogenetic tree, and metadata for amplicon dataset were uploaded into a Phyloseq object in R for further filtering and analysis [48]. ASVs corresponding to Chloroplasts, Mitochondria, Metazoa, and Embryophyceae were removed from the analysis. Additionally, ASVs were further filtered to include only those that were in the most abundant 99% of taxa in at least one of the samples in the dataset. Furthermore, any sample with less than 1000 reads was removed from the dataset. This stringent filtering left us with 46,423 ASVs across 286 samples. Bubble plots, alpha-diversity, and beta-diversity figures were constructed using ggplot2 and tidyverse packages [49]. Shannon-Weiner Indices and Aitchison distances were calculated from relative abundance-normalized data for alpha and beta-diversity analyses, respectively. A Global Wilcoxon Test using the rstatix package (version 0.7.2), with an adjusted p-value calculated using the “Holm” pairwise comparison, was conducted to determine whether Shannon diversity differed significantly. Analysis of similarities (ANOSIM) within vegan was used to test for differences in beta-diversity between families using 999 permutations [50]. Phylogenetic trees were constructed for select taxonomic groups of interest. To do this, first, the most abundant ASVs from the taxonomic group of interest were extracted from the filtered phyloseq object and aligned to a backbone alignment made up of sequences from PR2. Once the ASVs and references were properly aligned using HMMer, a RAxML (v8) maximum likelihood tree was constructed using the GTRCAT substitution model and 1,000 iterations [51]. Trees were uploaded and manipulated for publication in R using ggtree [52]. For comparisons across other published metabarcoding datasets, metaPR2 [53] was used to download the following datasets: OSD_2015_V4, OSD_2014_V4_LGC, and OSD_2014_V4_LW [54] representing the worldwide coastal sites and Malaspina_surface_2010_2011 [55], Tara_Ocean_V4 [56] representing the global open ocean. These were filtered of any metazoan or Embryophyceae ASVs before downloading. All bioinformatic scripts used in this study are available at https://github.com/delCampoLab/coral_euks.

Short-read amplicons were mined from NCBI’s short read archive (SRA) for corresponding published coral microbiome 16S rRNA gene data sequenced using the Illumina platform. SRA accessions were downloaded using the package “SRA tools”. CutAdapt was used to trim varying primer pairs covering differing hypervariable regions using generalized primers encompassing all variations in ambiguous bases to bulk cut primers of similar origin. Once trimmed, amplicon sequence variants (ASVs) were inferred using the DADA2 pipeline, and taxonomy was assigned using a combined reference database containing both Silva V138.1 [57] merged with the Coral Microbiome Database [58] with a redundancy threshold of 99% sequence similarity. A subset phyloseq object containing only V4 hypervariable region data was created using the R package “phyloseq” and a manually curated metadata table obtained from both SRA and information pulled from the publications. Community structure was analyzed using R package “microbiome” for alpha diversity metrics, and box plots were created using “ggplot2” to compare prokaryotic and eukaryotic community diversity index values.

## Results

### The diversity of the coral eukaryome

The diversity of the coral eukaryome is lower than the open ocean but higher than that of coastal environments (Fig. 1A). Compared to the coral microbiome’s bacterial component, the eukaryome generally exhibits greater diversity (Fig. 1B). However, this pattern varies across anthozoan families. For instance, the eukaryome diversity is particularly pronounced in Poritidae, Dendrophylliidae, and Acroporidae. In contrast, bacterial diversity is higher in Meandrinidae, Merulinidae, and Astrocoeniidae (Fig. 1C). When comparing the Shannon diversity of the Symbiodiniaceae versus the rest of the eukaryotes, we observe similar patterns across the anthozoan families, except for the Siderastreidae and the Actiniidae in which the patterns are inverted (Supplementary Fig. 1).

**Figure 1.**
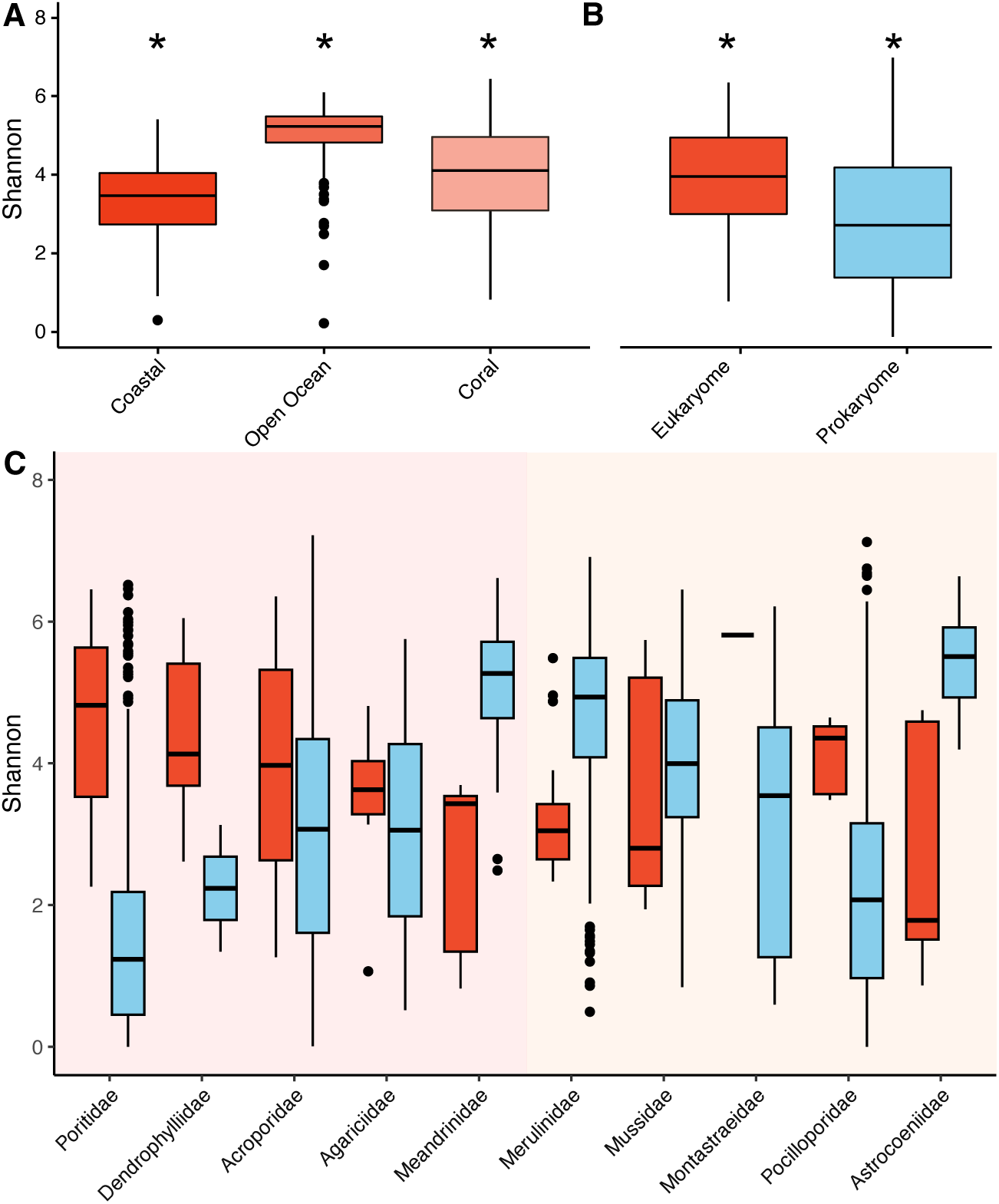
Alpha diversity (Shannon diversity index) across marine environments and coral microbiomes. **A.** Comparison of eukaryotic alpha diversity in three marine habitats: coastal waters (Ocean Sampling Day), open ocean (Tara Oceans), and coral-associated communities (this study). * All Shannon indices are significantly different from one another (Coastal to Coral p=7.86e-09, Open Ocean to Coral: p=4.21e-17, Open Ocean to Coastal p=1.07e-62, Global Wilcoxon Test). **B.** Alpha diversity of coral-associated eukaryotes (eukaryome - red) compared to prokaryotes (prokaryome - blue) across all samples. * Their Shannon indices are significantly different (prokaryome to eukaryome p=2.29e-14, Global Wilcoxon Test) **C.** Alpha diversity of coral-associated eukaryotes (red) and prokaryotes (blue) stratified by coral family. The shadowing groups the families in complexa (pink) and robusta (orange).

### The composition of the coral eukaryome

Not surprisingly the most common and abundant unicellular eukaryotes across most coral species are members of the Symbiodiniaceae family (Fig. 2A). Other prominent groups of unicellular eukaryotes include fungi, chlorophytes, ciliates, apicomplexans, other dinoflagellates distinct from Symbiodiniaceae, and rhodophytes (Fig. 2A). However, this overall pattern of relative abundance varies significantly across different coral families and groups. Among these, green and red algae exhibit exceptionally high abundances in scleractinian corals. Corallicolids are prevalent in robust corals, while Ichthyodinida and dinoflagellates are more abundant in Antipatharia (black corals) and octocorals. *Cladocopium* is the dominant Symbiodiniaceae symbiont in most coral families, except for the Meandrinidae, Mussidae, Monstastraeidae, and Gorgoniidae, which are dominated by *Breviolum,* and the Actiniidae, which are dominated by *Durusdinium*.

**Figure 2.**
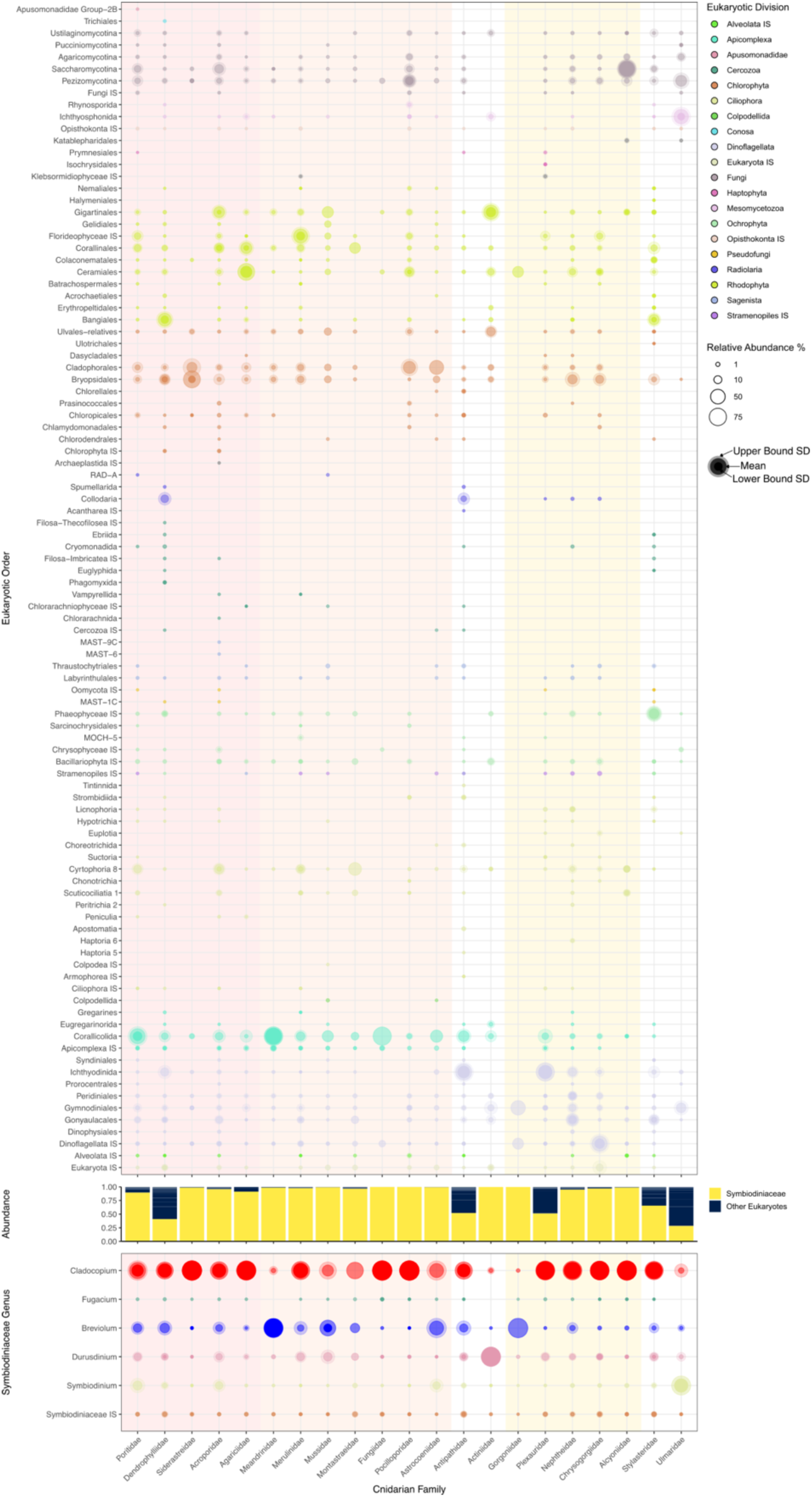
Microeukaryotic composition of healthy coral holobionts across families based on 18S rRNA metabarcoding data. Bubble plot depicting the taxonomic diversity and relative abundance of eukaryotic lineages detected in healthy coral samples, grouped by host cnidarian family on the x-axis. Taxa are organized by major eukaryotic orders (left) on the y-axis. Separating the bubble plots are bar graphs depicting the proportion of Symbiodiniaceae (yellow), compared to the remainder of microeukaryotes (dark grey). Symbiodiniaceae diversity is shown organized by genus (left). The shadowing groups the coral families in complexa (pink), robusta (orange), and octocorals (yellow).

**Figure 3.**
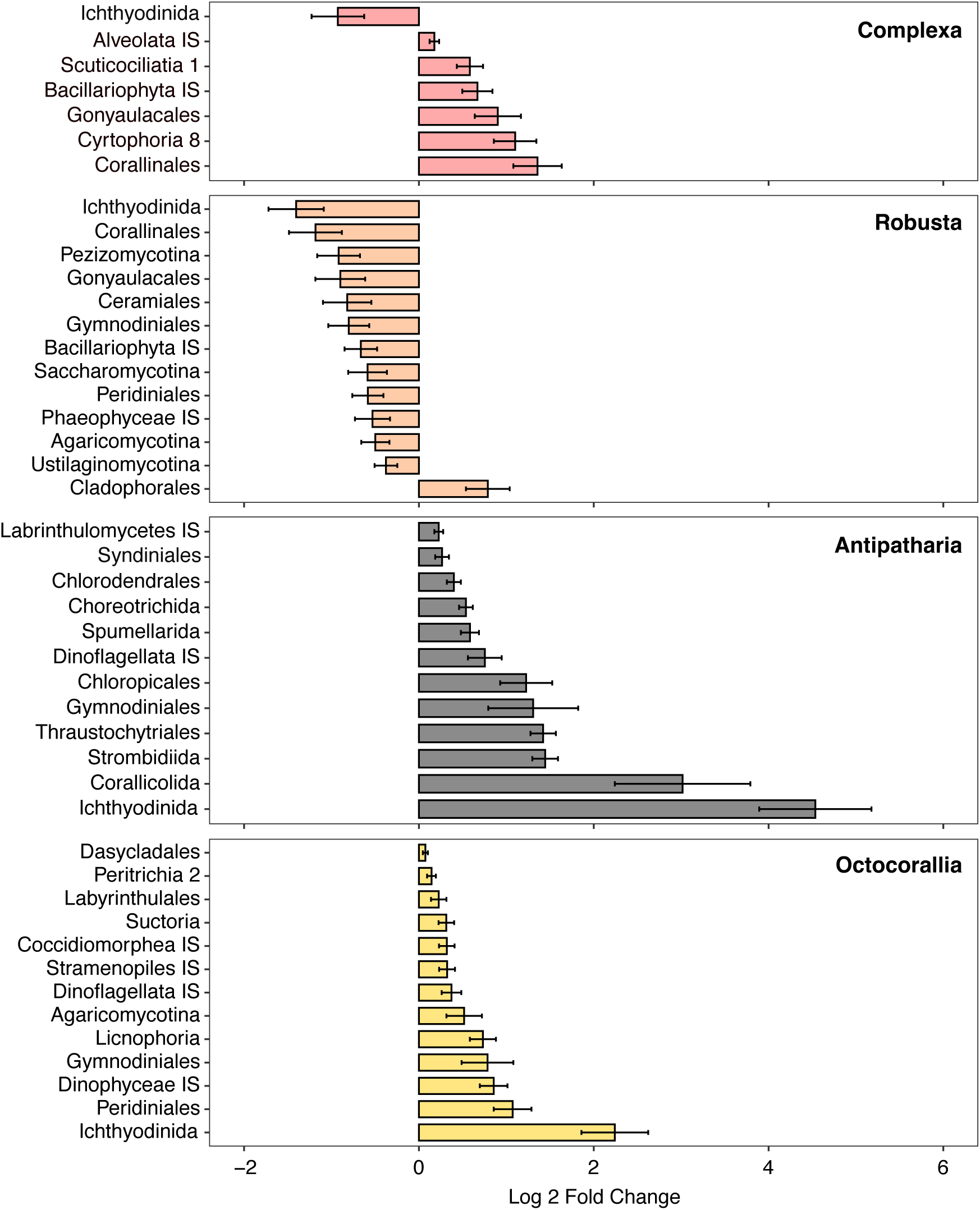
Differential abundance analysis of eukaryotic taxa across coral groups using ANCOM-BC. Log2 fold change in the relative abundance of eukaryotic orders across coral samples, as identified through ANCOM-BC. Positive values indicate enrichment, while negative values indicate depletion relative to the reference group. Only significantly differentially abundant taxa (adjusted *p* < 0.05) are shown.

The ANCOM analysis reveals that certain groups of anthozoans present differentially abundant groups of protists. Complexa scleractinian corals present significantly more Corallinales and more “Cyrtophoria 8” ciliates (close to twofold) than the rest of the corals, at the same time, complexa corals present significantly less Ichthyodinida. Robusta scleractinian corals present significantly more Cladophorales than the rest of the analyzed corals (2 times), while presenting fewer Corallinales and Ichthyodinida (2 times). Antipatharians present significantly more Ichthyodinida (4 times) and Corallicolids (3 times). Octocorals present more Ichthyodinida (2 times) and more dinoflagellates other than Symbiodiniaceae (Peridiniales, Gymnodiniales, and IS Dinophyceae).

In diseased corals, there is a generalized shift in the community composition across the coral families (Supplementary Fig. 2A), and in certain families, shifts in specific dominant communities are evident. For example, in Siderastreiade the Bryopsidales disappear completely in diseased corals, and there is generally a shift from *Cladocopium* to *Breviolum* and an increase in the presence of Gymnodiniales. We also observed a dramatic shift in diseased Pocilloporidae, from *Cladocopium* to *Durusdinium*. In other groups, such as the Acroporidae from Japan, there is a clear divergence between the eukaryome of healthy and diseased corals (Supplementary Fig. 2B), but too few samples to outline specific recurring trends.

Some widespread groups, such as Corallicolida and Symbiodiniaceae, exhibit distinct biogeographical distributions, suggesting environmental or host-specific influences on their diversity and composition (Figure 4).

**Figure 4.**
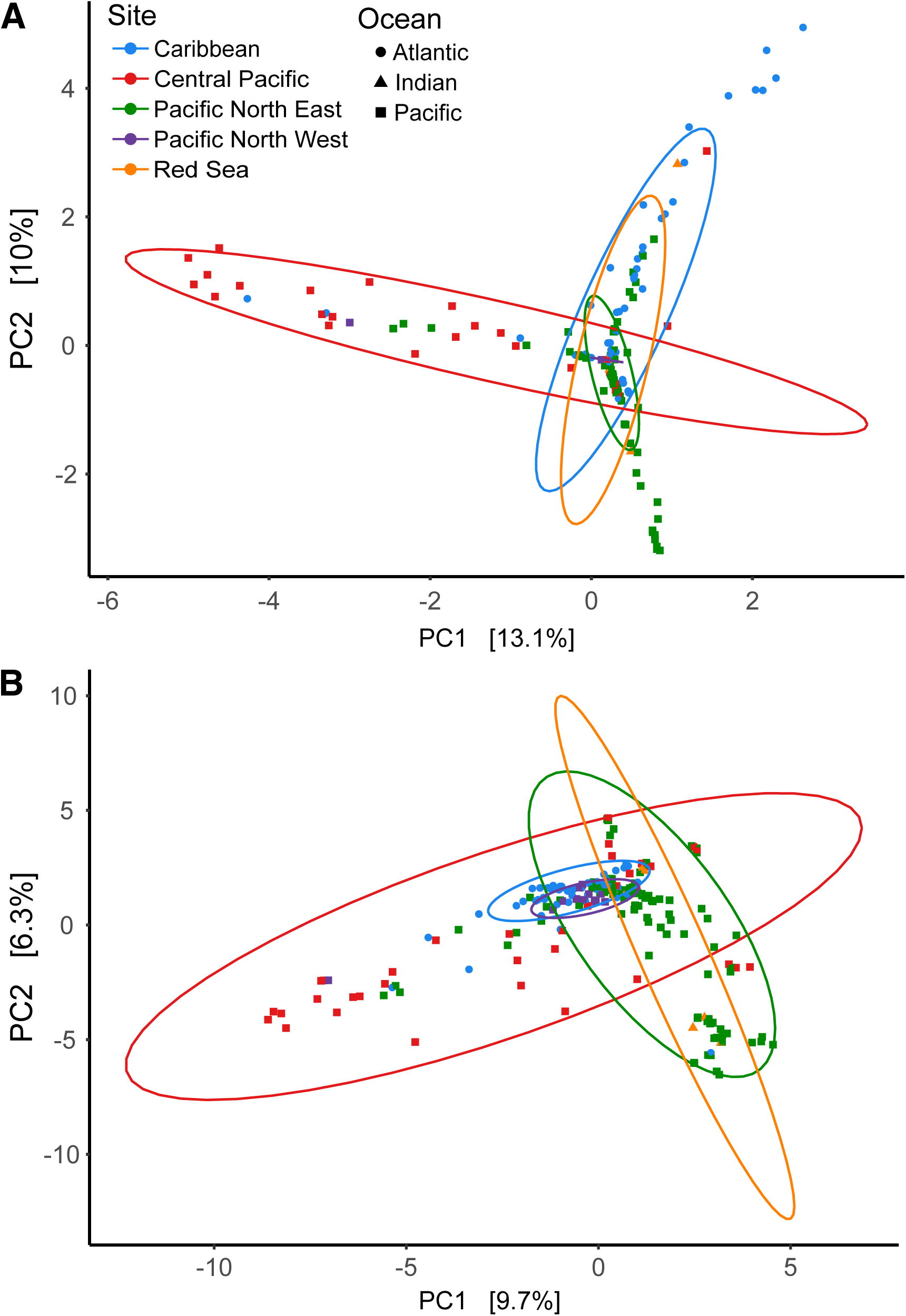
Biogeographic structuring of key eukaryotic symbiotic lineages across ocean basins. **A.** Principal component analysis (PCA) of *Corallicolida* community composition across sampling sites. Samples are color-coded by site. **B.** PCA of *Symbiodiniaceae* community composition across sampling sites. Samples are color-coded by site.

### Main Protist Groups in Corals

Corallicolida are present across all analyzed coral groups, including scleractinians, black corals, and octocorals (Figure 5A). Identified hosts include members of the Stylasteridae, which is an important range expansion for the group, which have otherwise only been observed to infect anthozoans. Some ASVs, such as ASV_401, ASV_373, ASV_369, ASV_584, ASV_290, ASV_296, and ASV_349, are abundant but are not always the most widespread. Of the described ASVs, only ASV_4962 is related to a known species (*Corallicola aquarius*). Specific patterns emerge among coral groups: ASV_3877, ASV_2299, and Groups 1 and 4 dominate in hexacorals, while ASV_290 and Group 5 dominate in octocorals. Interestingly, we found no ASVs among the analyzed corals assigned to *Anthozoaphila gnarlus,* which has been repeatedly microscopically identified and isolated from Caribbean hosts over many years, perhaps due to different biases in the two approaches.

**Figure 5.**
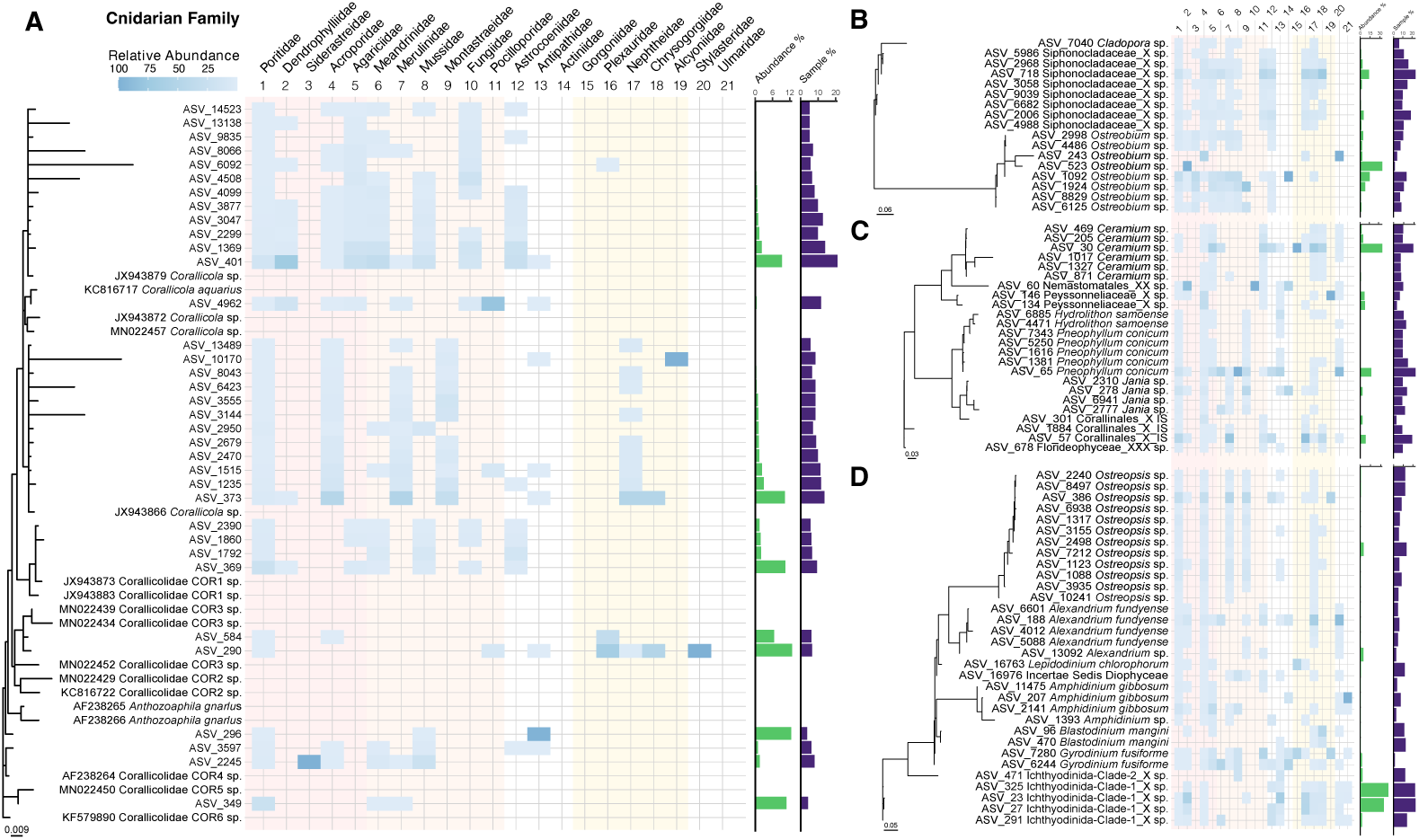
Phylogenetic placement and taxonomic distribution of major symbiotic eukaryotic lineages in corals. ASVs classified into each of the analyzed groups were EPA-placed onto a backbone 18S rRNA gene maximum likelihood phylogenetic tree for each group. The relative abundance of each ASV is shown in the heatmap across cnidarian families. The green histogram on the right side of each panel shows the relative abundance of each ASV within the analyzed group, and the purple histogram its prevalence across all the analyzed samples. **A.** *Corallicolida* (Apicomplexa). **B.** Ulvophyte green algae (Clorophyta, primarily *Ostreobium* and related siphonous lineages). **C.** Red algae taxa (Rhodophyta). **D.** Dinoflagellates (Dinoflagellata, excluding Suessiales). The shadowing groups the coral families in complexa (pink), robusta (orange), and octocorals (yellow).

*Ostreobium* and Siphonocladales are the most abundant and widespread chlorophytes (Fig. 5B), with Ostreobium predominantly associated with hexacorals and Siphonocladales also present in octocorals. Within the latter, ASV_718 is particularly abundant in octocorals. *Ostreobium* exhibits high diversity, with some ASVs broadly distributed across coral families, while others, such as ASV_523, show host specificity, being found exclusively in Siderastreidae. In parallel, the most abundant and widespread rhodophytes include *Ceramium*, *Pneophyllum*, and an unidentified member of the Corallinales (ASV_57) (Fig. 5C), all of which occur in both hexacorals and octocorals. Among these, ASV_30 (*Ceramium*) and ASV_65 (*Pneophyllum*) are the most dominant ASVs. Turning to dinoflagellates, the genus *Ostreopsis* displays the highest diversity, although it is not especially abundant (Fig. 5D), while Ichthyodinida emerges as the most abundant group within the dinoflagellates, particularly in octocorals, with ASV_23 and ASV_27 being its most prominent representatives. Among ciliates, ASVs assigned to *Mirodysteria decora*, especially ASV_130, are both abundant and prevalent across coral samples (Supplementary Fig. 3A). Finally, despite the limited taxonomic resolution provided by the short 18S fragment, fungi are confirmed as integral components of the coral eukaryome (Fig. 2, Supplementary Fig. 3B), with ASV_101, assigned to *Debaryomyces hansenii*, standing out as both abundant and widespread

The Symbiodiniaceae genus with the highest diversity is *Cladocopium*, followed by *Brevioulum*, *Durusdinium*, and *Symbiodinium*. Most individual ASVs are present in relatively low abundance and are distributed differently across anthozoan families. However, a handful of ASVs represent most of the reads and are present in virtually all samples; two correspond to *Cladocopium* (ASV_1 and ASV_2) and one to *Breviolum* (ASV_3), with a few less abundant ASVs assigned to *Durusdinium* and *Symbiodinium*. We identified several hotspots of unrecognized diversity in Symbiodiniaceae: one sister to *Fugacium* and *Cladocopium,* one sister to *Cladocopium*, and one at the base of Symbiodiniaceae. No ASVs related to *Effrenium* or *Fugacium* were detected in our dataset (Fig. 6).

**Figure 6.**
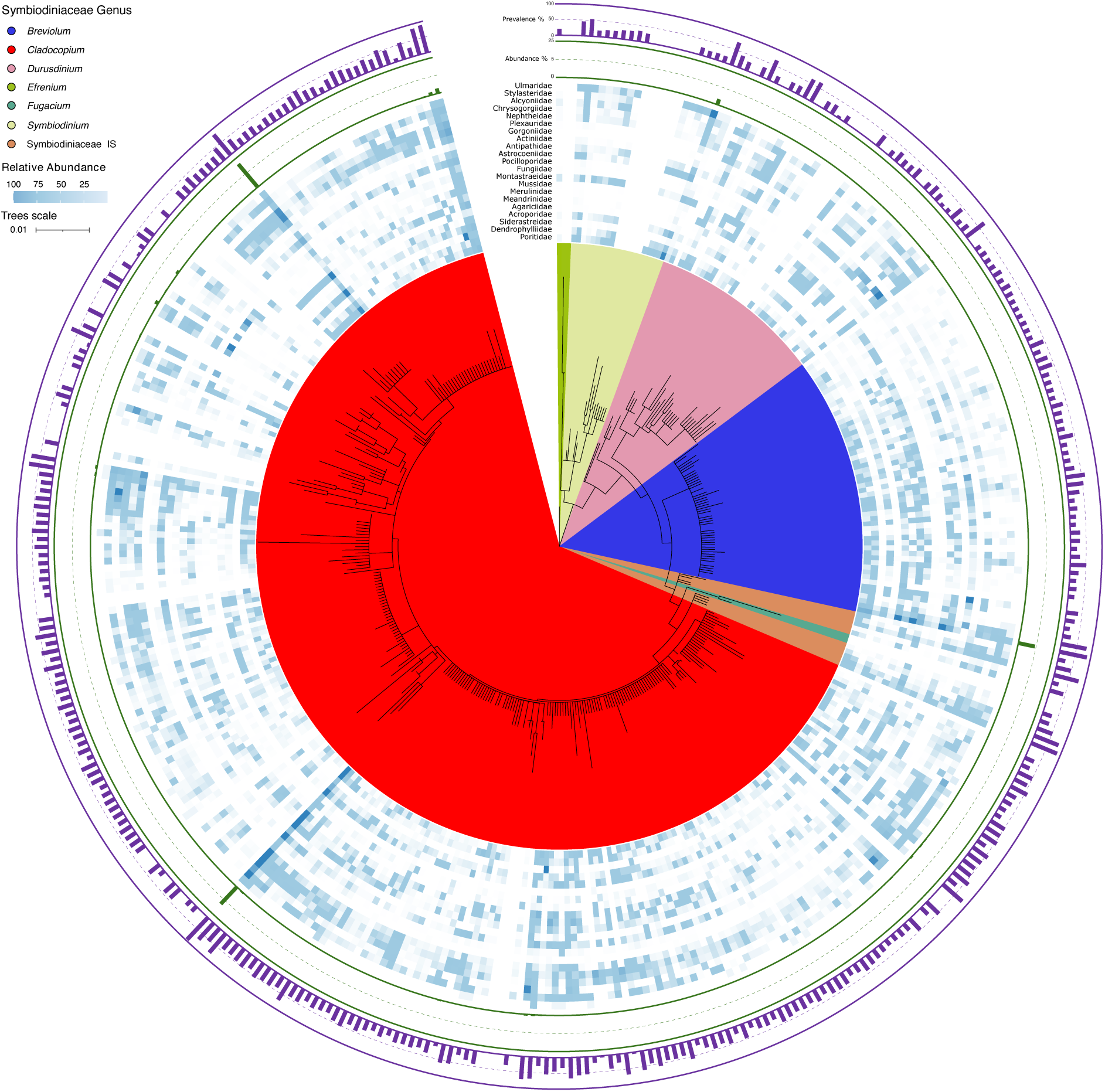
Phylogenetic placement and taxonomic distribution of Symbiodiniaceae in corals. ASVs classified as Suessiales were EPA-placed onto a backbone 18S rRNA gene maximum likelihood phylogenetic tree for the group. The relative abundance of each ASV is shown in the heatmap across cnidarian families. The green histogram on top of the heatmap shows the relative abundance of each ASV within the analyzed group, and the purple histogram its prevalence across all the analyzed samples.

## Discussion

These analyses cover a significant diversity of anthozoans and other cnidarians from multiple sites worldwide. Accordingly, they expand the described diversity of coral-associated protists by a factor of three and reveal eukaryotic microbial diversity comparable to that of the open ocean (Fig. 2). Indeed, the coral eukaryome is more diverse than the free-living eukaryotic communities of some marine environments, such as the coast, and more surprisingly is even more diverse than the prokaryome of some coral groups (Fig. 1A).

Across coral species, the most common and abundant unicellular eukaryotes are not surprisingly members of the Symbiodiniaceae family, consistent with their well-established role as key symbionts in corals [7]. The Symbiodiniaceae are, of course, well-studied in corals; however, the 18S rRNA gene is not typically the preferred marker choice for Symbiodiniaceae metabarcoding, with the most common one being the ITS (Internal Transcribed Spacer) [59]. The 18S rRNA gene marker nonetheless revealed a great diversity of Symbiodiniaceae, although interestingly, only a handful of ASVs were found to be both abundant and widespread. There are several possible reasons for this: (1) amplification biases are different for the primers used here, (2) only a handful of species that we can associate with the abundant and widespread ASVs dominate the pool of Symbiodiniaceae sampled here, or (3) that the Symbiodiniaceae contain many copies of the rRNA operon and what we observe is variability in 18S rRNA gene diversity rather than species diversity.

Dinoflagellates beyond Symbiodiniaceae are also widespread across all corals, with the families Gymnodiniales and Gonyulacales being particularly abundant in octocorals, together with a significant number of ASVs with an uncertain position within the Dinophyceae. *Ostreopsis* (Gonyaulacales), a diverse and widespread genus in our dataset, is a potentially toxic dinoflagellate [60], as is *Alexandrium fundyense* [61], which we also find associated with corals. Some of these relationships have been reported before in coral reefs [62], and may either be due to them being eaten by the corals, as they are common inhabitants of the coral reef waters, or because they are playing an undefined role.

Green and red algae are both particularly abundant and present in scleractinian corals. Some of these have already been identified as epibionts [63] and endoliths [64, 65] of coral reef substrates. In the case of green algae, we observe that Cladophorales and Bryopsidales are the two most abundant and widespread orders linked to *Cladophora sp.* [66] and *Ostreobium quekettii* [17], respectively. We also find green algae associated with octocorals; in this case, ASVs are identified as Siphonocladales, which are common algae in intertidal and subtidal habitats [67]. Within red algae, we observe relatively high diversity across multiple orders, including a novel group within Florideophyceae. Considering the abundant presence of red algae in coral reefs, we contemplate two possibilities: (1) these are not part of the coral holobiont and were instead from the surroundings (despite each sample being carefully rinsed), or (2) certain life phases of the red algae are linked to the corals as epibionts or endoliths, such as the conchocelis phase of some *Porphyra* species and other Bangiales (Laborel and Le Campion-Alsumard, 1979; Tribollet et al., 2018).

The widespread presence of Corallicolida has been reported previously; here, we confirm that they are abundant in several families of Scleractinia and are significantly overrepresented in black corals. In the case of black corals, this correlates with a high abundance of Ichthyodinida, which are also abundant in octocorals. These two groups have been shown to negatively correlate in octocorals, where their presence is linked to heat-stress sensitivity [37]. With both groups being widespread across corals, they may play similar roles in several coral families where they co-occur, including scleractinians. In the case of the corallicolids, we also observe that particular ASVs dominate in different coral families. For instance, the ASVs related to *Corallicola aquarius* and Corallicolidae COR1 are associated with scleractinian corals, while ASVs associated with COR3 are dominant in octocorals and hydroids. This suggests a certain degree of coevolution between the Corallicolida and their hosts. Still, they have previously been argued to move rapidly between hosts, putatively through a vector [68], suggesting a complex interplay between the two may be true.

Corallicollids also display differential geographic distribution in our data (Figure 4) which has been proposed based on a smaller sample size of full-length 18S rRNA gene [69]. This suggests that correlations between historical colonization events and larval dispersion of corals with the geographical patterns of Corallicolida may exist. We also report the presence of Corallicolida in *Stylaster* sp. and *Stylanteca* sp. (Stylasteridae, Hydrozoa), representing the first record of this symbiont in Cnidaria outside the Anthozoa.

Ciliates, well-known coral-associated protists commonly linked to disease [70], are also widespread across the corals in our data, particularly *Mirodysteria decora* (Cyrtophoria), which is also relatively abundant (Supplementary Fig. 3A). Little is known about the biology of *M. decora,* apart from the fact that they are marine commensal ciliates [71]. However, based on what is known about ciliates in corals more generally, it seems likely that they live as epibionts, feeding on coral-associated microbes and gastric cavity debris [72]. Another ciliate that is widespread but not abundant across all anthozoan families is *Philaster apodigitiformis*. This ciliate has been identified as the causative agent of the recent mass mortality of the Caribbean Sea urchin, *Diadema antillarum* (Hewson et al., 2023). Considering we found these scuticociliates associated with healthy corals, they are most likely commensals, which could act as reservoirs for the sea urchin parasite.

Lastly, we find a wide variety of fungi, and despite the 18S rRNA gene not being the best marker for this group, we can still identify significant fungal diversity in our data. The most abundant and widespread coral fungi ASVs correspond to *Debaryomyces hansenii*, a marine yeast of interest to the biomedical and biotechnological industries for its antimicrobial capacity [73], which has been shown to enhance the defensive capabilities of fish skin (Sanahuja et al., 2023). Other common fungi reported in marine environments, such as *Aspergillus* [74] or *Malassezia restricta* [68], are also present in the anthozoan eukaryome.

Consistent with the expectation that parasite richness should be greater on pristine reefs where host life cycles remain intact, we found that putatively parasitic protists were widespread and often abundant across coral groups in remote, relatively undisturbed sites. Corallicolida (Apicomplexa), Ichthyodinida, and several ciliates (e.g., *Mirodysteria decora*) were detected in nearly all coral families examined, with particularly high relative abundance in octocorals and black corals. Importantly, these taxa were not restricted to diseased or degraded colonies, but were also common in apparently healthy corals, supporting the view that diverse parasite communities are a baseline component of the holobiont in intact ecosystems. By contrast, shifts in the eukaryome associated with disease were not characterized by simple gains or losses of parasite groups, but by changes in their relative dominance within the community. For instance, in diseased Pocilloporidae, *Cladocopium* was replaced by *Durusdinium*, while in Siderastreidae, Bryopsidales disappeared entirely and Gymnodiniales increased. These results suggest that while parasite richness is maintained across reef conditions, disease states reflect reorganization of existing symbiotic and parasitic lineages rather than invasion by novel taxa.

Our global survey shows that microbial eukaryotes are consistent and widespread components of the coral holobiont. Groups such as Symbiodiniaceae, Corallicolida, Ostreobium, Ichthyodinida, and ciliates were recovered across coral families and ocean basins, often in both healthy and diseased colonies. Their ubiquity suggests that these associations are not incidental but reflect long-term evolutionary relationships.

Symbiodiniaceae diversity, for example, was extensive, yet dominated by a few globally persistent ASVs, consistent with the conservation of stable partners over deep time. Likewise, the broad occurrence of Corallicolida across anthozoans supports an ancient association, possibly tracing evolutionary transitions from phototrophy to parasitism within coral hosts.

At the same time, our data reveal evidence of recurrent colonization and turnover. Ostreobium exhibited host-specific ASVs, while groups such as Ichthyodinida and non-Symbiodiniaceae dinoflagellates showed strong biases toward particular coral clades, indicating repeated episodes of specialization. Importantly, disease was associated not with the emergence of novel eukaryotes, but with shifts in dominance among resident lineages, such as the replacement of *Cladocopium* by *Durusdinium*. Taken together, these findings support the view that corals, among the oldest extant animals, have acted as both refugia and selective environments for protist evolution. The coral holobiont thus represents not only a present-day ecological consortium but also a long-term evolutionary archive of animal–protist symbiosis.

## Conclusions

The technical challenge of using metabarcoding to examine the microbial eukaryotes associated with eukaryotic (animal) hosts is a substantial barrier that has prevented a comprehensive understanding of the role of protists and fungi on coral reef ecology. With this study, we show one way to overcome this challenge and describe the first taxonomically broad approximation of the diversity of microbial eukaryotes associated with corals. Along with revealing new associations and correlations, these results more generally highlight the importance of studying all the microbes within the holobiont, and that protists associated with corals do play key roles in the ecosystem that require further investigation from both coral reef scientists and protistologists.

## Declarations

### Ethics approval and consent to participate

Not applicable.

### Consent for publication

Not applicable.

## Availability of data and material

The 18S rRNA amplicon reads are deposited in the NCBI Sequence Read Archive (PRJNA482746 and PRJNA1373245).

## Author’s contribution

JdC, FR, and PJK conceived and designed the study. JdC, BK, AA, MDF, KW, MV, and PJK collected and provided samples for the study. JdC, AMB, and BAW analyzed the metabarcoding data. JdC, AMB, and PJK wrote the manuscript; all the co-authors contributed with reviews, comments, and discussion.

## Acknowledgements

We would also like to thank the Frost Institute for Data Science & Computing and the Centro de Supercomputación de Galicia for providing the supercomputing resources necessary for this project.

## Supplementary Figures

**Supplementary Figure 1.**
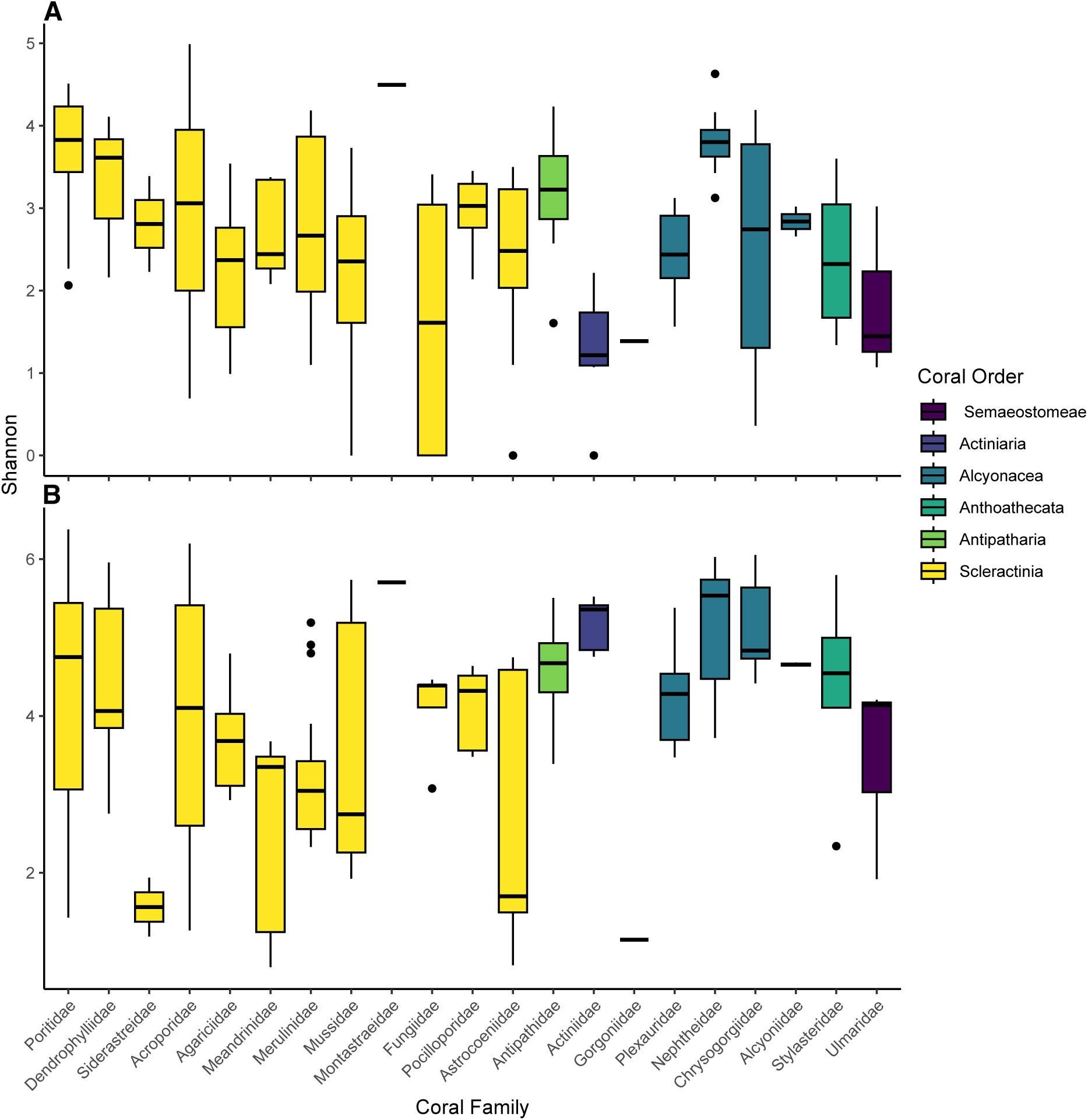
Alpha diversity (Shannon diversity index) of coral-associated eukaryotes by host taxonomy. **A.** Alpha diversity of all non-*Symbiodiniaceae* eukaryotic lineages associated with coral samples, stratified by coral family. **B.** Alpha diversity of *Symbiodiniaceae* across coral families.

**Supplementary Figure 2.**
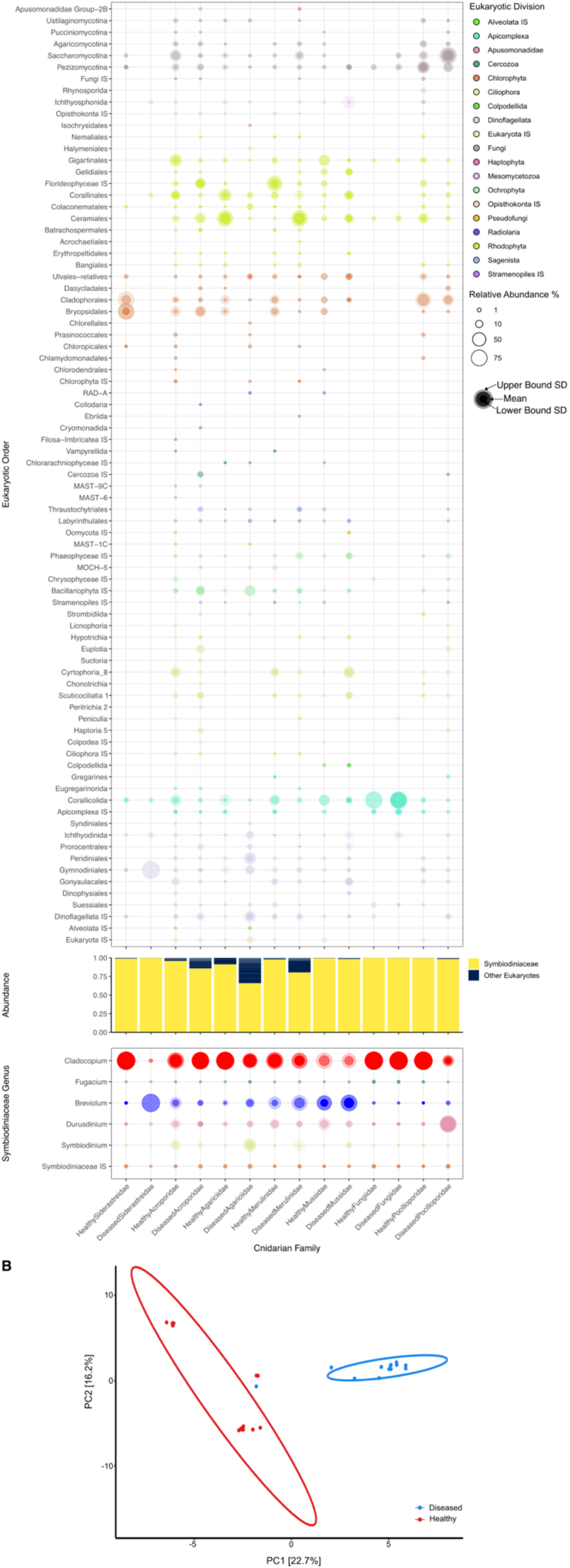
A. Microeukaryotic composition of healthy vs diseased coral holobionts across families based on 18S rRNA metabarcoding data. Bubble plot depicting the taxonomic diversity and relative abundance of eukaryotic lineages detected in healthy and diseased coral samples, grouped by host cnidarian family. Taxa are organized by major eukaryotic orders (left). Symbiodiniaceae diversity is shown organized by genus (left). The shadowing groups the coral families in complexa (pink) and robusta (orange).**B. Structuring of key eukaryotic symbiotic lineages across disease states.** Principal coordinates analysis (PCoA) of *Acroporidae* samples from Japan, comparing healthy and diseased individuals.

**Supplementary Figure 3.**
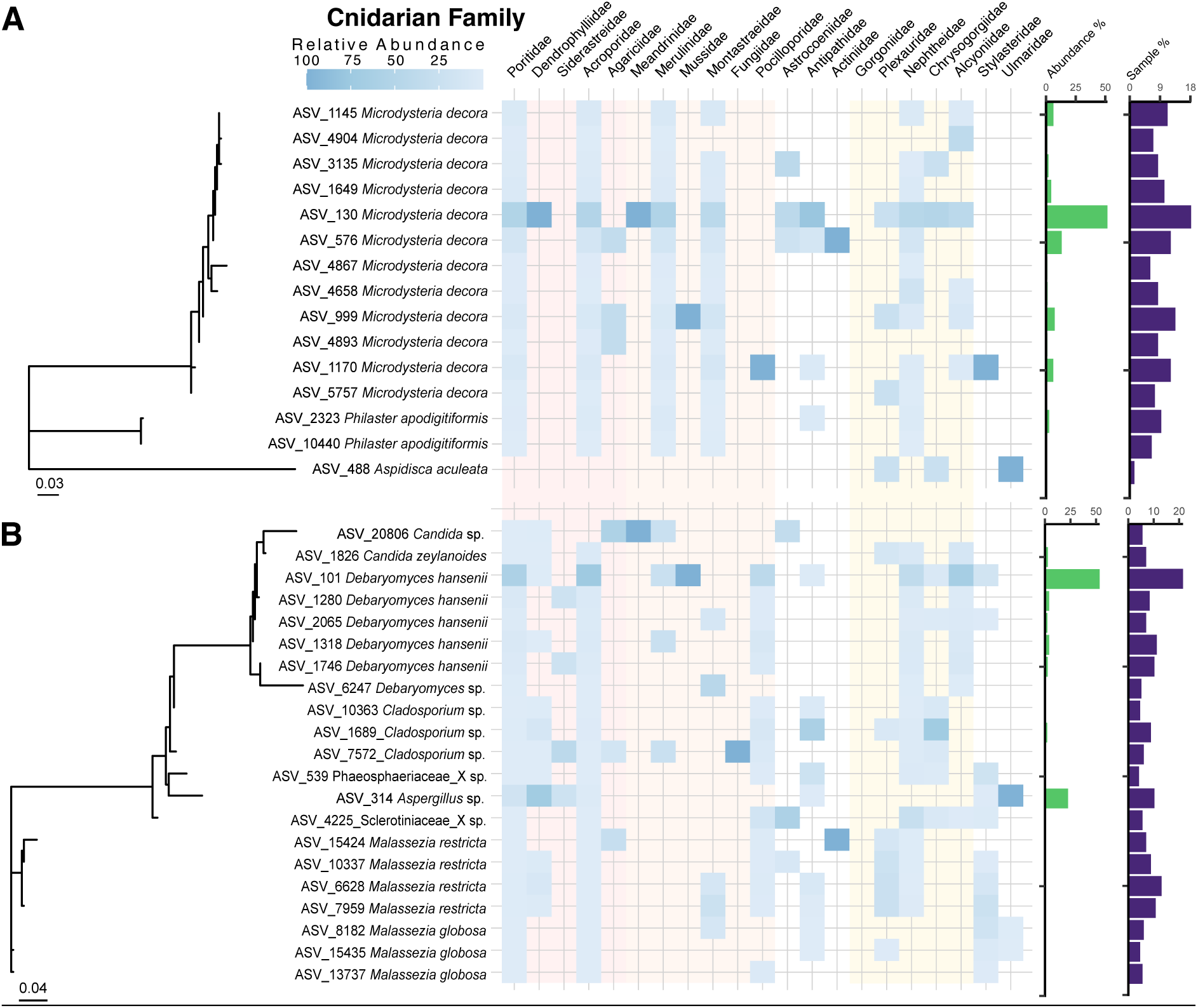
Phylogenetic placement and taxonomic distribution of ciliate and fungal ASVs in corals. The most abundant ASVs classified into fungi (**A**) and ciliates (**B**) were EPA-placed onto a backbone 18S rRNA gene maximum likelihood phylogenetic tree for each group. The relative abundance of each ASV is shown in the heatmap across cnidarian families. The green histogram on the right side of each panel shows the relative abundance of each ASV within the analyzed group, and the purple histogram its prevalence across all the analyzed samples. The shadowing groups the coral families in complexa (pink), robusta (orange), and octocorals (yellow).

**Supplementary Table 1. Sampled coral species by site and taxonomic classification.** Comprehensive list of all coral specimens included in this study, organized by sampling site and annotated with complete taxonomic classification.

