## Supplementary figures and images for "Global diversity and distribution of coral-associated protists"

### Supplementary Figure 1

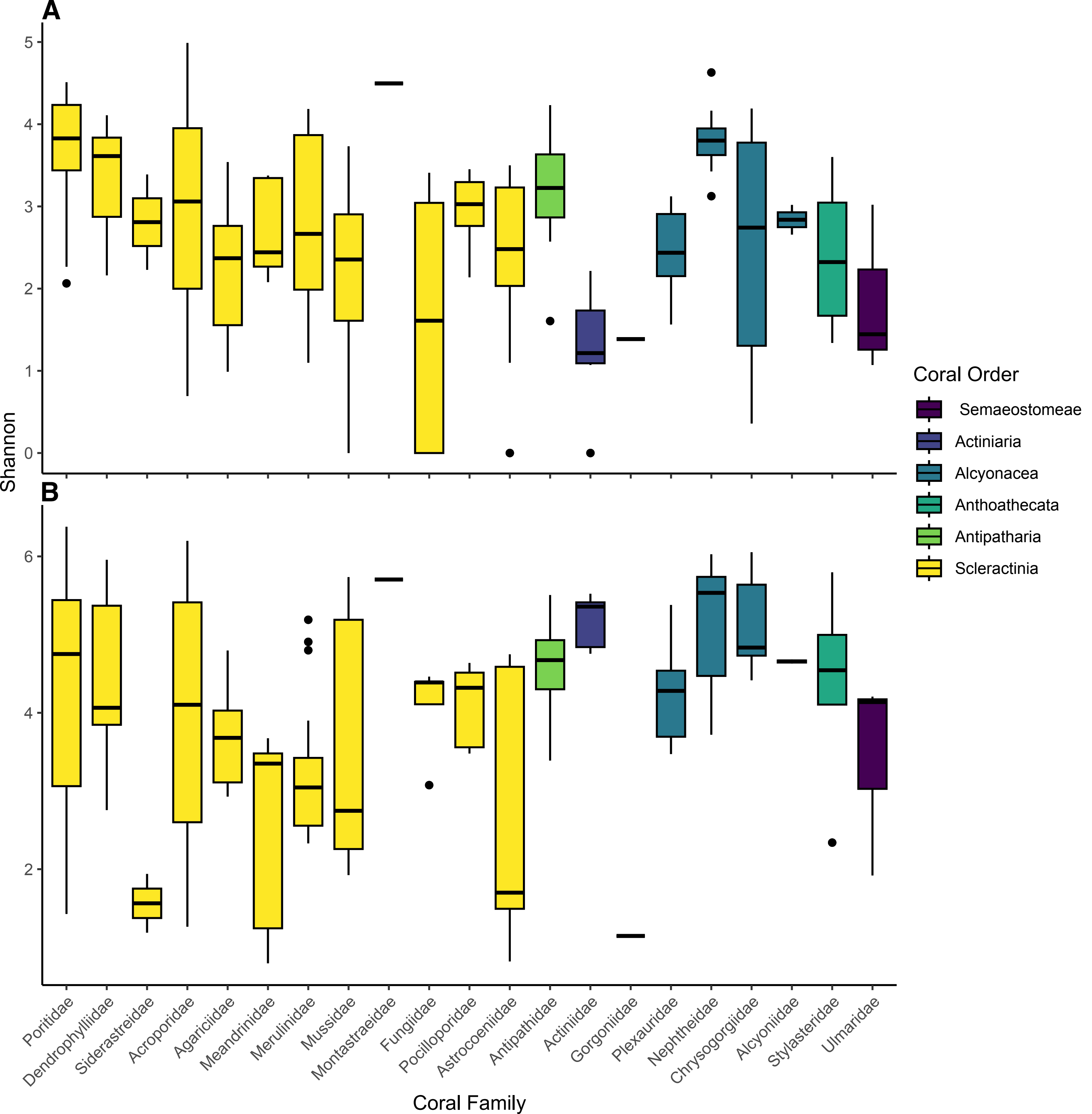

### Supplementary Figure 2

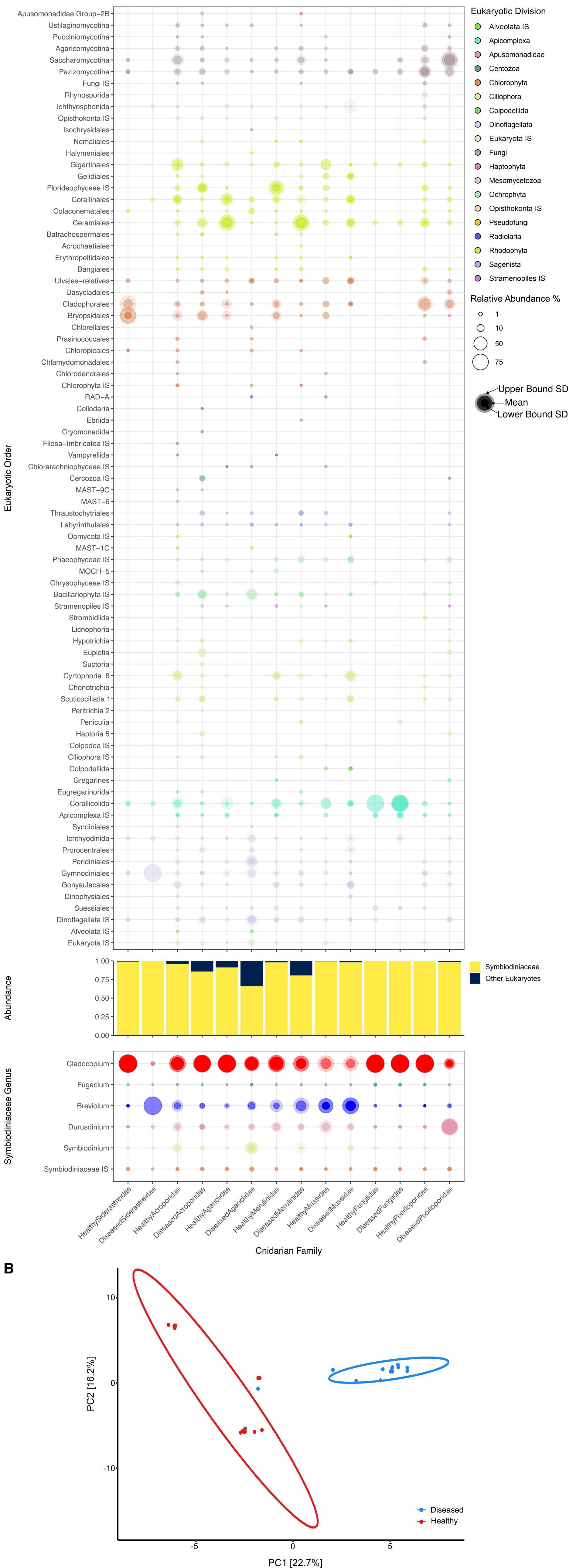

### Supplementary Figure 3

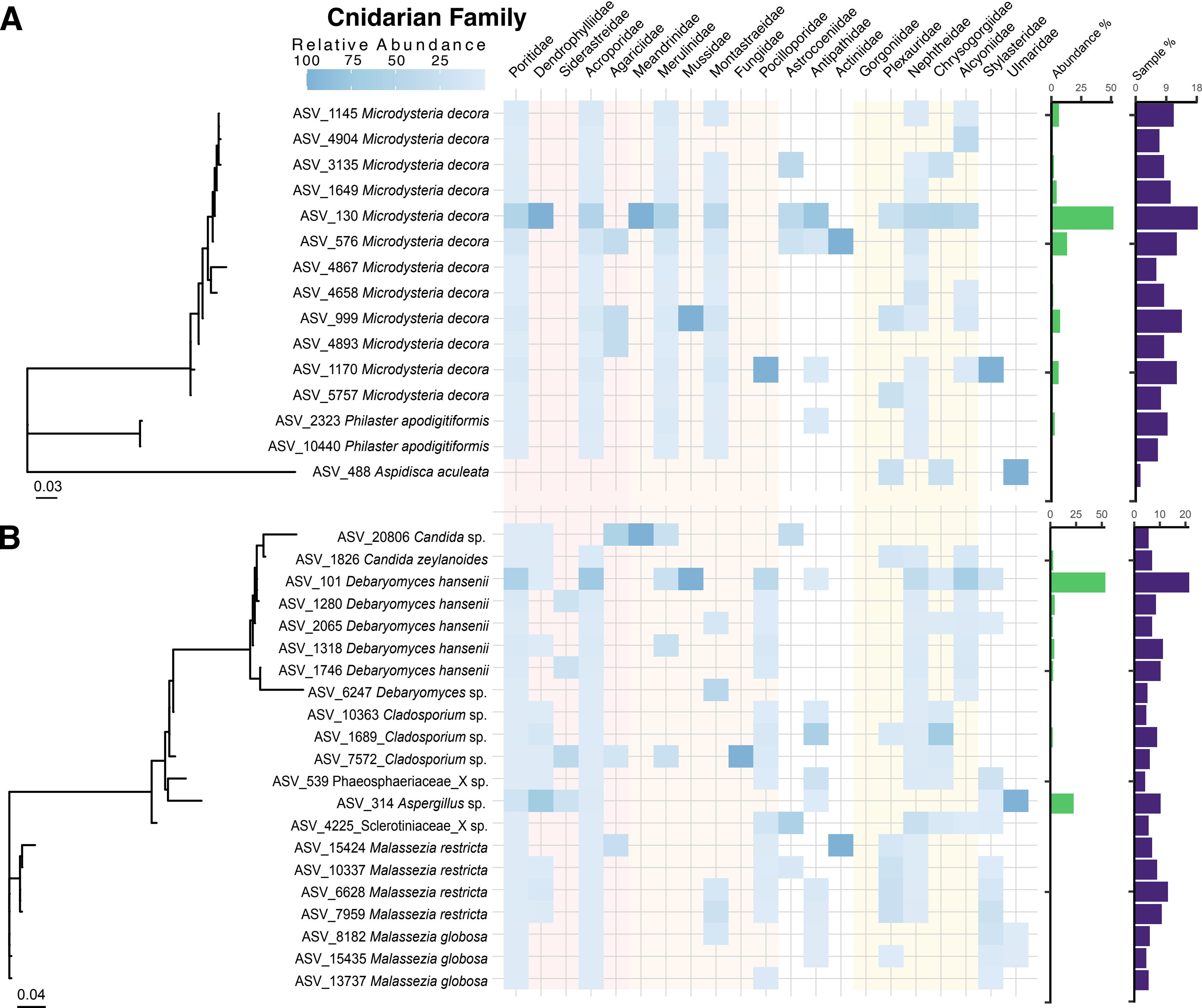
